# Suppressed Osteocyte Perilacunar / Canalicular Remodeling Plays a Causal Role in Osteoarthritis

**DOI:** 10.1101/534768

**Authors:** Courtney M. Mazur, Jonathon J. Woo, Cristal S. Yee, Aaron J. Fields, Claire Acevedo, Karsyn N. Bailey, Tristan W. Fowler, Jeffrey C. Lotz, Alexis Dang, Alfred C. Kuo, Thomas P. Vail, Tamara Alliston

## Abstract

Osteoarthritis (OA), long considered a primary disorder of articular cartilage, is commonly associated with subchondral bone sclerosis. However, the cellular mechanisms responsible for changes to subchondral bone in OA, and the extent to which these changes are drivers of or a secondary reaction to cartilage degeneration, remain unclear. In knee joints from human patients with end-stage OA, we found evidence of profound defects in osteocyte function. Suppression of osteocyte perilacunar/canalicular remodeling (PLR) was most severe in OA subchondral bone, with lower protease expression, diminished canalicular networks, and disorganized and hypermineralized extracellular matrix. To determine if PLR suppression plays a causal role in OA, we ablated the PLR enzyme MMP13 in osteocytes, while leaving chondrocytic MMP13 intact. Not only did osteocytic MMP13 deficiency suppress PLR in cortical and subchondral bone, but it also compromised cartilage. Even in the absence of injury, this osteocyte-intrinsic PLR defect was sufficient to reduce cartilage proteoglycan content and increase the incidence of cartilage lesions, consistent with early OA. Thus, in humans and mice, osteocyte PLR is a critical regulator of cartilage homeostasis. Together, these findings implicate osteocytes in bone-cartilage crosstalk in the joint and identify the causal role of suppressed perilacunar/canalicular remodeling in osteoarthritis.

## INTRODUCTION

Osteoarthritis (OA), the most common chronic joint disease, is a leading cause of pain and disability worldwide (1). OA irreversibly damages articular cartilage and the surrounding tissues, compromising joint function and mobility of over 30 million Americans (2). Abundant research efforts have investigated cartilage and its interactions with other joint tissues in order to understand the underlying mechanisms of OA initiation and progression (3–5). Still, no permanent disease-modifying therapies exist short of joint replacement.

A major unanswered question in the field is the extent to which subchondral bone plays a causal role in the pathogenesis of OA. Subchondral bone sclerosis, together with progressive cartilage degradation, is widely considered a hallmark of OA (3,6). Though much correlative evidence indicates the coordinated degradation of subchondral bone and cartilage, causality is difficult to ascertain, because analyses are often conducted on tissues with end-stage disease or in models in which both the bone and cartilage are affected. Therefore, the cellular mechanisms responsible for OA-related changes in subchondral bone, and the extent to which these changes are drivers of or a secondary reaction to cartilage degeneration, remain unclear. In particular, little is known about the role of osteocytes.

Recent reports have reinvigorated interest in the active role of osteocytes in remodeling their surrounding bone matrix – a process called perilacunar/canalicular remodeling (PLR) (7–10). PLR is a dynamic process by which osteocytes secrete matrix metalloproteinases (MMPs) (11–13), cathepsin K (CatK) (8), and other enzymes (14,15) to dynamically resorb and then replace the local bone matrix. PLR maintains bone material properties (11,16,17), systemic mineral homeostasis (8,10), and the canalicular channels that facilitate osteocyte communication, mechanosensation, and nourishment (7,18,19). Several known regulators of bone homeostasis, including TGF-β (17), SOST (15), parathyroid hormone (8,20), and Vitamin D (21,22), regulate PLR to support the metabolic and mechanical function of the skeleton. Although PLR is a fundamental mechanism by which osteocytes maintain bone homeostasis, its role in the maintenance of subchondral bone and the progression of joint disease remain unclear.

To elucidate the role of PLR in disease, we previously investigated osteonecrosis, a progressive and severe joint disease in which subchondral bone mechanically fails with painful collapse of the articular surface (23,24). We found that glucocorticoids, a major risk factor associated with osteonecrosis (23), suppress PLR and cause the same changes in subchondral bone of mice as seen in glucocorticoid-induced human osteonecrosis (25). Although suppression of PLR is clearly associated with the degradation of subchondral bone in osteonecrosis, it was not possible to isolate the effects of osteocytes from the systemic influence of glucocorticoids. Given that the health of articular cartilage depends upon subchondral bone for mechanical and vascular support (26,27), we hypothesized that signs of PLR dysregulation may also accompany the much more common joint disease, osteoarthritis. We tested this hypothesis by examining specific hallmarks of PLR suppression and their relationship to cartilage degeneration in OA of the human knee. To further isolate the contributions of PLR to joint disease, we evaluated the bone and joint phenotypes of a novel mouse model with ablation of MMP13 from osteocytes but not chondrocytes. Our results implicate a novel, causal role for osteocyte PLR in the pathogenesis of OA.

## RESULTS

### Degeneration of the osteocyte lacunocanalicular network in human osteoarthritis

To determine if osteocytic PLR is affected by osteoarthritis (OA) in humans, we compared subchondral bone in tibial plateaus from patients with end-stage OA to that of cadaveric donors with no clinical evidence of joint disease. As expected, tibial plateaus from patients with OA had gross degeneration of the articular cartilage (Supp. Fig. 1A-F) and radiographic evidence of subchondral bone sclerosis, particularly on the medial side of the joint (Supp. Fig. 1G-I). Histological analysis confirmed the cartilage degeneration and subchondral bone sclerosis in OA specimens (Fig. 1A-F). Consistent with prior reports (6), both the subchondral bone plate and trabeculae were thicker in OA (Fig. 1B, E, F) than in controls (Fig. 1A, C, D). Relative to the cadaveric controls, OA tibial plateaus showed decreased cartilage thickness, reduced Safranin-O positive proteoglycan staining in the superficial zone, and cartilage fibrillation (Fig. 1C-F), resulting in significantly higher OARSI scores (Supp. Fig. 1B). Although the control cartilage was more intact than the OA cartilage overall, the medial compartment showed more evidence of degeneration in both OA and control specimens. Therefore, the subsequent analyses of subchondral bone compared defined regions of interest (Fig. 1A, B) between the control and OA specimens, as well as between the medial and lateral side of the same specimen.

**Figure 1:**
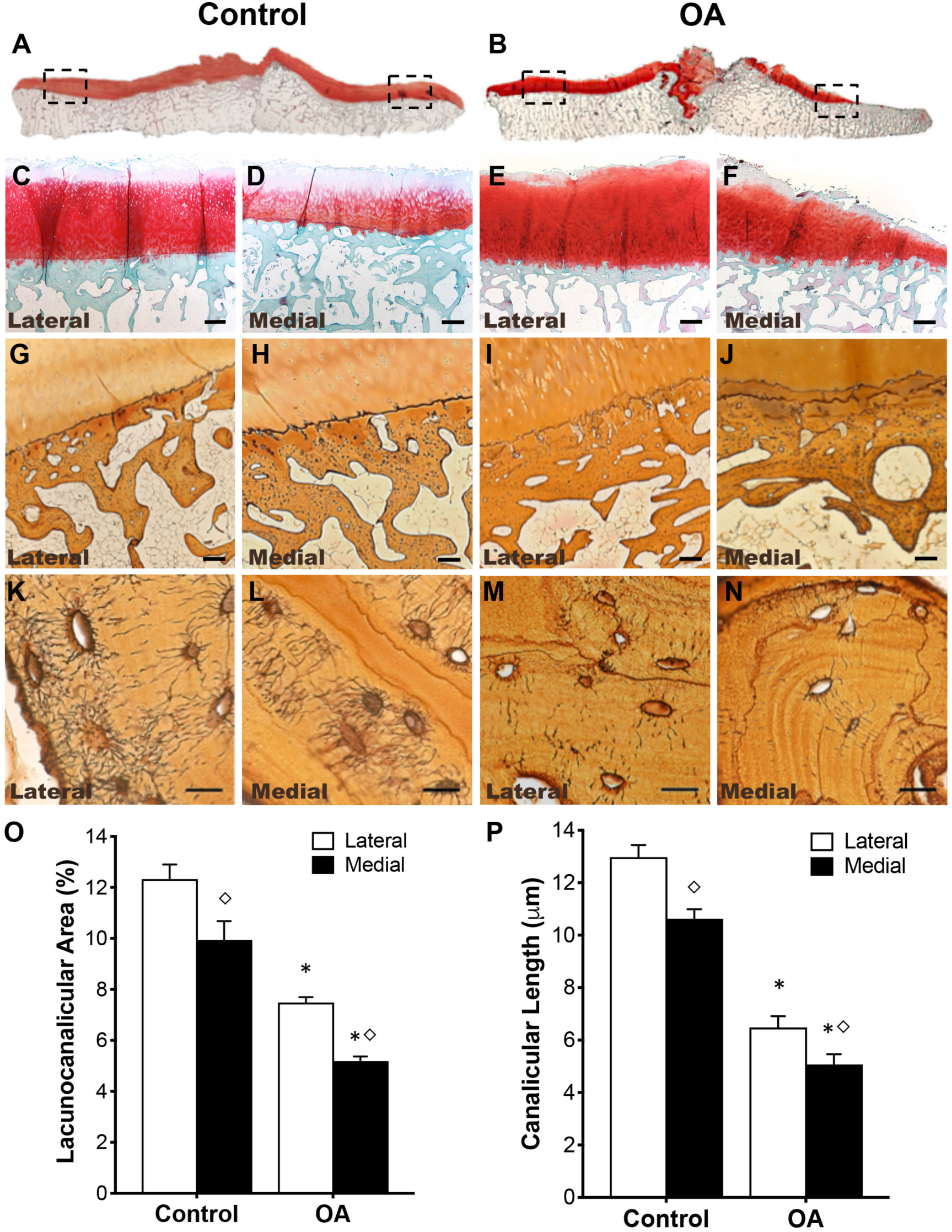
Disrupted lacunocanalicular networks in human OA subchondral bone. Control cadaveric (A, C, D) and OA (B, E, F) specimens stained with Safranin-O/ Fast Green and imaged at 0.5x (A, B) or 10x (C-F) magnification displayed differences in articular cartilage and subchondral bone morphology on the lateral and medial sides of the tibial plateau. Subsequent analyses compared the indicated regions of interest (black boxes in A, B) between control and OA specimens, and between the less affected lateral side with the more severely degraded medial side. These identified regions of interest in Ploton silver stained sections were evaluated at low (4x, G-J) and high (100x, K-N) magnification to visualize the lacunocanalicular network of subchondral bone. Quantification of lacunocanalicular area normalized to bone area (O) and canalicular length (P) revealed significant OA-dependent reductions in both parameters (n=5). Scale bars are 400 µm in C-F, 200 µm in G-J, and 20 µm in K-N. *p<0.05 compared to respective regions of control specimens, ◊p<0.05 between regions by Holm-Sidak post-hoc tests.

While OA-dependent differences in osteoblast and osteoclast function have been described (3,6), the effect of OA on osteocytes is not well-defined. Therefore, we evaluated osteocyte PLR in subchondral bone by studying one of its key hallmarks, the lacunocanalicular network (LCN). Silver staining revealed that cadaveric control subchondral bone (Fig. 1G, H, K, L) had both more abundant and apparently longer canalicular projections than the subchondral bone from OA patients (Fig. 1I, J, M, N). The dramatic degeneration of the canalicular network in OA bone was particularly evident in the medial side of the tibial plateau, where cartilage degeneration was most severe (Fig. 1N). Quantitative analysis revealed significant OA-dependent reductions in the total osteocyte lacunocanalicular area (38-46%) and canalicular length (51-54%) relative to cadaveric controls, consistent with this hallmark feature of PLR suppression (8,11–13,17,25,28) (Fig. 1O, P). Therefore, the reduced canalicular length and lacunocanalicular area in OA subchondral bone strongly suggests that PLR is suppressed in OA.

### Collagen disorganization and hypermineralization in human OA subchondral bone

In mouse models of PLR suppression and in human osteonecrotic subchondral bone, loss of lacunocanalicular area is often accompanied by collagen disorganization and hypermineralization of the bone extracellular matrix (ECM) (11,25,28). Therefore, we evaluated the organic and mineral constituents of OA subchondral bone. Collagen fibrils in OA subchondral bone showed qualitatively less birefringence relative to the control tissue (Fig. 2A-H). Upon quantification, collagen linearity was significantly lower in OA specimens compared to control specimens on both the lateral and medial sides. Furthermore, collagen was significantly less aligned on the medial side of the joint than on the lateral side in both groups (Fig. 2I).

**Figure 2:**
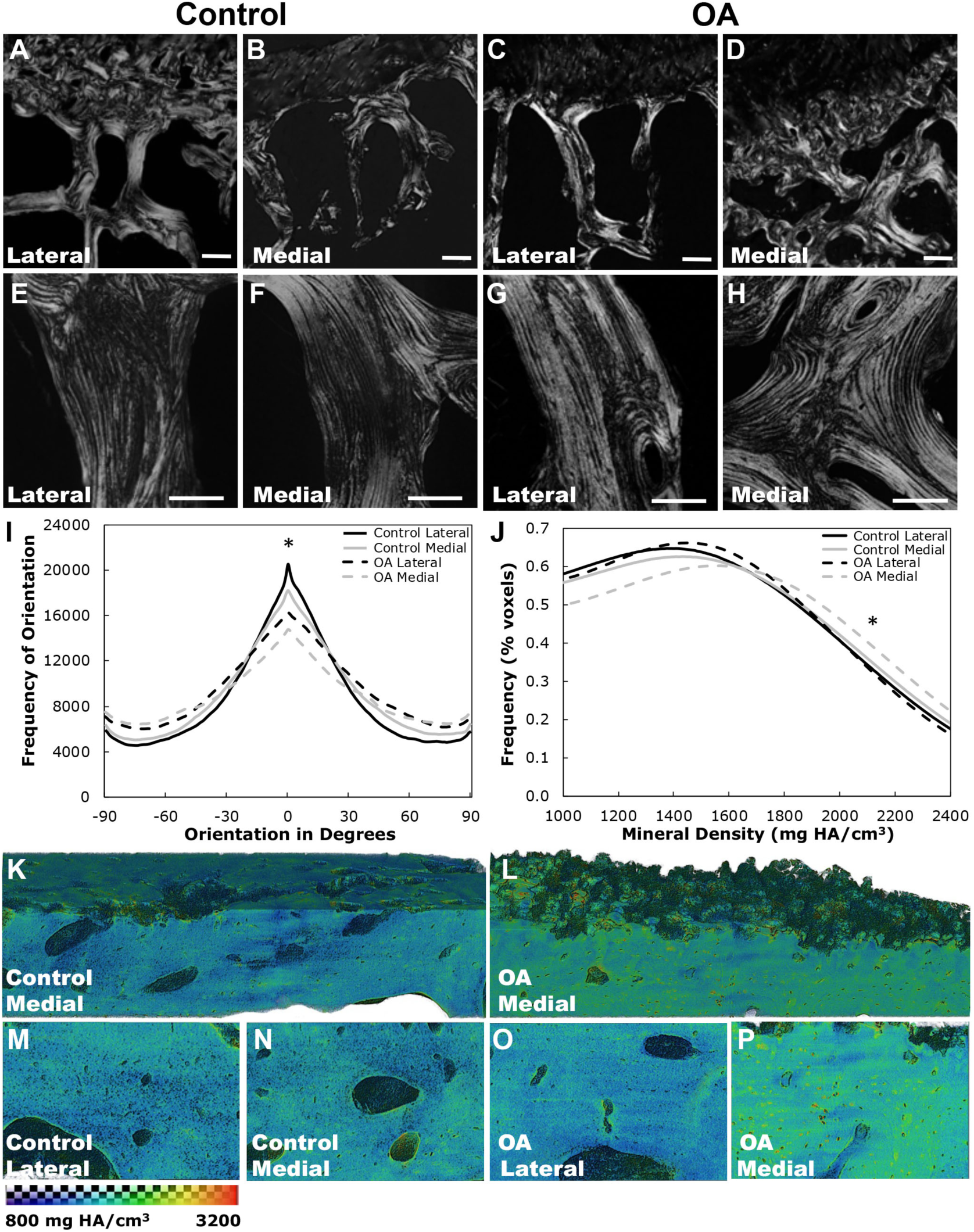
Collagen disorganization and hypermineralization in human OA subchondral bone. Control cadaveric (A, B, E, F, n=5) and OA (C, D, G, H, lateral n=5, medial n=4) specimens stained with Picrosirius Red and imaged at low (4x, A-D) or high (40x, E-H) magnification using polarized light microscopy revealed differences in subchondral bone collagen organization. Collagen is less organized in OA samples than control samples and in medial regions than lateral regions. The distribution of collagen orientation shows significant differences between all four groups (I). Synchrotron Radiation X-ray micro-computed tomography (SRµT) of subchondral bone from the lateral (M, O) and medial (K, L, N, P) sides of cadaveric control (K, M, N) and OA (L, O, P) tibial plateaus showed a qualitative increase in mineralization, according to the colorimetric scale (800 to 3200 mg HA/cm^3^), in 3D reconstructed (K, L) and 2D high magnification (M-P) images. The distribution of mineralization through the subchondral bone from each region (J) confirms the shift in the peak mineralization level in OA medial specimens (n=5) compared to OA lateral (n=4) and each control region (n=2). Scale bars are 200 µm in A-D and 100 µm in E-H. *p<0.05 by mixed model with random intercepts.

Consistent with these site- and disease-dependent patterns, hypermineralization of subchondral bone was most pronounced in specimens from the medial side of the OA tibial plateau. Medial OA specimens also portrayed a rougher subchondral surface than control specimens (Fig. 2K, L), and a significant shift in the distribution of mineral density relative to any of the other sites (Fig. 2J, M-P). Therefore, OA is accompanied by subchondral bone collagen disorganization and bone matrix hypermineralization, concordant with suppressed PLR (11,25,28).

### Reduced osteocyte expression of PLR enzymes in human OA subchondral bone

Given that OA subchondral bone shows multiple signs of suppressed PLR, we sought to evaluate the expression of key enzymes implicated in PLR by osteocytes. Immunohistochemistry (IHC) revealed qualitatively lower levels of MMP13 (Fig. 3A-H) and Cathepsin K (CatK, Fig. 3J-Q) protein expression in subchondral bone of the medial OA tibial plateau relative to healthy controls and relative to the less severely affected lateral OA tibial plateau. Accordingly, the percentage of MMP13-positive osteocytes was lower in OA subchondral bone by 20% in medial portions and 10% in lateral portions relative to their respective control sites (Fig. 3I). The percentage of CatK-positive osteocytes was 24% lower on the medial side and 13% lower on the lateral side in OA subchondral bone compared to relative cadaveric controls (Fig. 3R). Interestingly, the percentage of MMP13-positive osteocytes is strongly correlated with lacunocanalicular area and with canalicular length for each sample and region (Fig. 3S-T).

**Figure 3:**
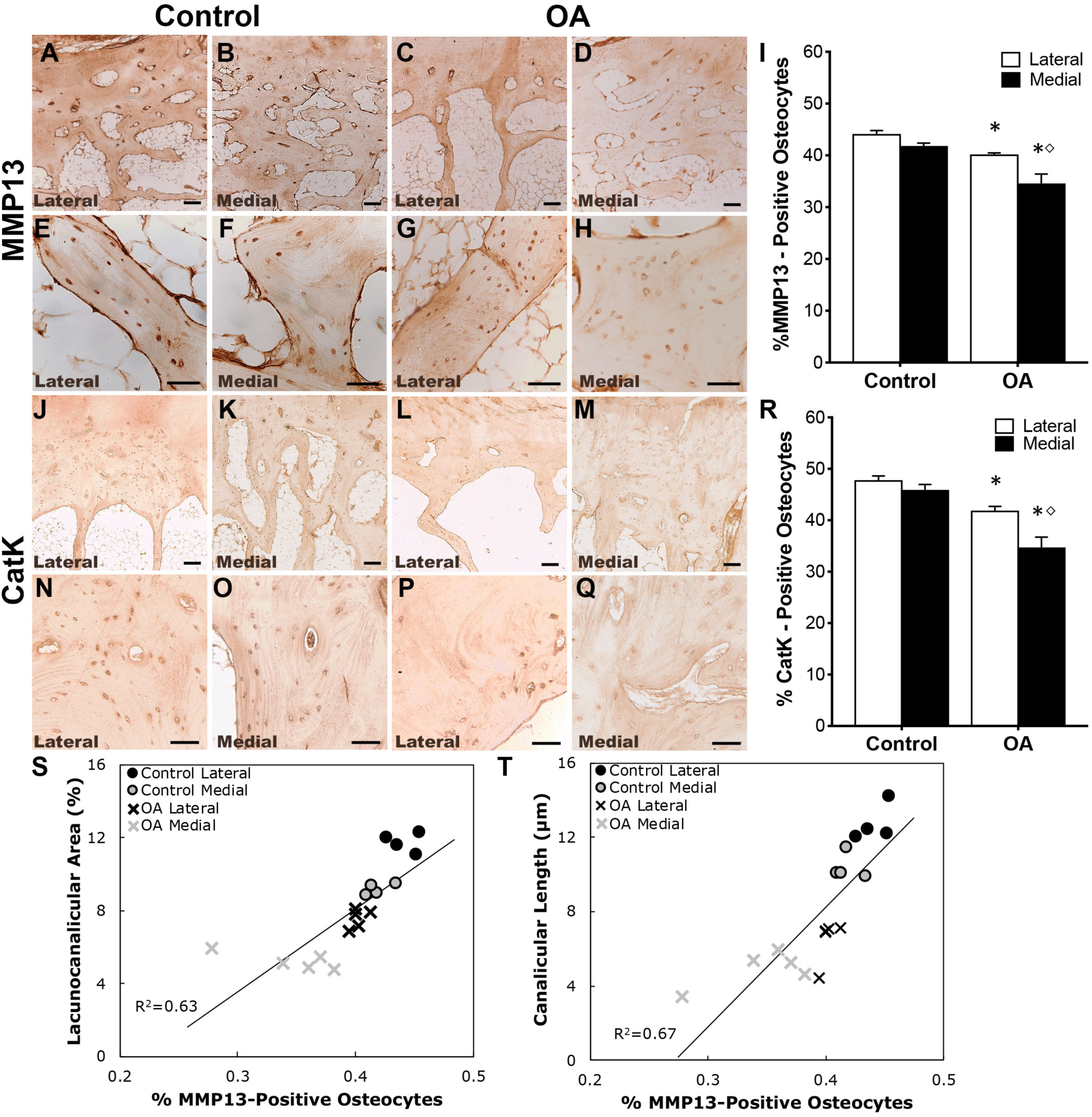
Suppressed PLR enzyme expression in human OA subchondral bone. Immunohistochemical analysis of MMP13 (A-H) and Cathepsin K (CatK, J-Q) levels and localization in subchondral bone from control and OA tibial plateau specimens was performed at low (4x, A-D, J-M) and high (20x, E-H, N-Q) magnification. Qualitative and quantitative analyses show diminished MMP13 and CatK expression in the OA tibial plateau, with a significant reduction in the percentage of osteocytes stained positively for MMP13 (I, control n=4, OA n=5) and CatK (R, n=5) in both regions. Furthermore, Pearson’s product-moment correlation indicates that percent of MMP13-positive osteocytes is strongly correlated with lacunocanalicular area (S, r=0.79, p<0.0001) and canalicular length (T, r=0.82, p<0.0001). Scale bars are 100 µm. *p<0.05 compared to respective regions of control specimens, ◊p<0.05 between regions by Holm-Sidak post-hoc tests.

Together these results suggest that human OA is correlated with PLR suppression in subchondral bone, as demonstrated by repression of key PLR enzymes in subchondral bone, loss of lacunocanalicular area, collagen disorganization, and hypermineralization. To further investigate whether PLR plays a causal role in joint disease, we generated mice with a targeted deletion of MMP13 from osteocytes. We previously reported that systemic ablation of MMP13 suppresses PLR (11), however, chondrocyte expression of MMP13 contributes to cartilage degradation (29), and ablation of MMP13 in chondrocytes is chondroprotective (30,31).

Therefore, we characterized the bone and joint phenotypes of mice with a novel, osteocyte-intrinsic ablation of MMP13.

### Specific ablation of MMP13 expression in osteocytes

An established floxed MMP13 allele (32) was deleted under control of DMP1-Cre (9.6-kb promoter) (33), resulting in mice with a targeted deletion of MMP13 in osteocytes. DMP1-Cre^+/-^;MMP13^fl/fl^ (MMP13^ocy-/-^) animals are born at the same rate and are grossly similar to their DMP1-Cre^-/-^;MMP13^fl/fl^ (wildtype) littermates, with no significant differences in weight or lifespan.

We validated the tissue-specific reduction in MMP13 expression at the transcriptional and translational level. In femoral cortical bone of MMP13^ocy-/-^ animals, immunofluorescence revealed 37% fewer MMP13-positive osteocytes, while expression in periosteal and endosteal cells was maintained (Fig. 4A-G). This result was corroborated by 63% reduced MMP13 mRNA expression in humeri cleaned of marrow and periosteum (Fig. 4H). Importantly, the expression pattern of MMP13 in growth plate and articular cartilage chondrocytes was unchanged in MMP13^ocy-/-^ mice (Supp. Fig. 2E-G, J-L). Therefore, the MMP13^ocy-/-^ mouse model is appropriate to observe differences in bone and joint phenotypes arising from changes in osteocyte-derived MMP13.

**Figure 4:**
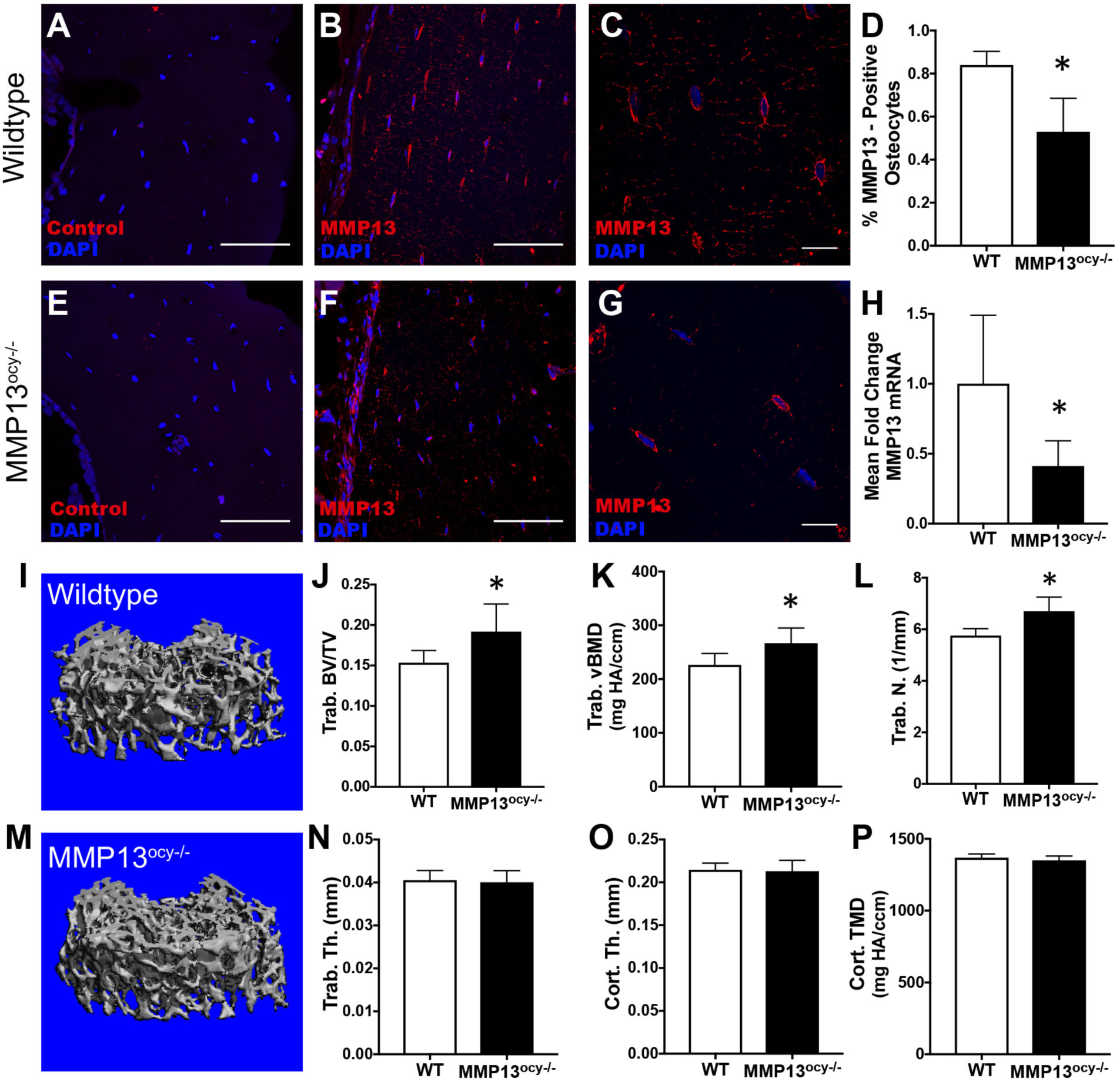
Osteocyte-specific reduction in MMP13 expression causes increased trabecular bone mass. In cortical bone, the number of osteocytes stained positive for MMP13 is reduced in MMP13^ocy-/-^ bone (F, G, D, n=9) compared to wildtype (B, C, n=8). No differences were seen in negative control images (A, E). MMP13 mRNA was also significantly reduced in MMP13^ocy-/-^ bone (H, n=9) compared to wildtype (n=7). µCT of distal femoral trabecular bone (I, M, n=11) shows that bone volume fraction (J) and volumetric bone mineral density (K) is increased in MMP13^ocy-/-^ bone due to an increase in trabecular number (L) rather than trabecular thickness (N). Cortical bone, however, did not present differences in thickness (O) or mineral density (P). Scale bars are 10 µm for C and G and 50 µm elsewhere. *p<0.05 between genotypes by unpaired t-test.

Trabecular bone volume is increased in mice with systemic ablation of MMP13 and in other models of PLR suppression (11,17,32). To determine if deletion of osteocyte-intrinsic MMP13 is sufficient to alter bone mass, we used µCT to analyze trabecular and cortical bone mass and microarchitecture. Relative to wildtype mice, MMP13^ocy-/-^ femurs had a 25% increase in trabecular bone volume fraction (Fig. 4I, J, M) due to a 16% increase in trabecular number (Fig. 4L) and a corresponding decrease in trabecular spacing (Supp. Fig. 2A) with no change in trabecular thickness (Fig. 4N). MMP13^ocy-/-^ femurs also show an increase in volumetric bone mineral density (Fig. 4K) and a decrease in SMI reflecting a shift to more plate-like microarchitecture (Supp. Fig. 2B). The mRNA levels or ratio of RANKL and OPG mRNA expression do not account for these differences (data not shown). Cortical bone thickness and total mineral density were normal in MMP13^ocy-/-^ femurs (Fig. 2O-P). Therefore osteocyte-intrinsic MMP13 is sufficient to alter trabecular bone volume and mineralization, and we suspect that the PLR phenotype might be affected.

### Suppressed PLR in MMP13^ocy-/-^ bone

To determine the role of osteocyte-intrinsic MMP13 in PLR, we evaluated the osteocyte LCN and collagen alignment, both of which are sensitive to PLR suppression, including in mice with systemic ablation of MMP13 (11). The LCN of femoral cortical bone is visibly disrupted by osteocyte-intrinsic MMP13 deficiency (Fig. 5A, B, E, F). Canalicular length in MMP13^ocy-/-^ mice is reduced by 20% (Fig. 5I) with no significant change in lacunar area (Fig. 5K) or lacunar density (data not shown). This decrease in canalicular length occurs in a coordinated manner across the medial, lateral, anterior and posterior regions of MMP13^ocy-/-^ cortical bone (Fig. 5I). We consistently observed a small but significant decrease in peak collagen alignment in MMP13^ocy-/-^ bones compared to wildtype bone in the anterior region (Fig. 5C, G, J). In the other regions studied (Fig. 5D, H), no differences in collagen linearity were detected despite the change in PLR activity suggested by LCN analysis.

**Figure 5:**
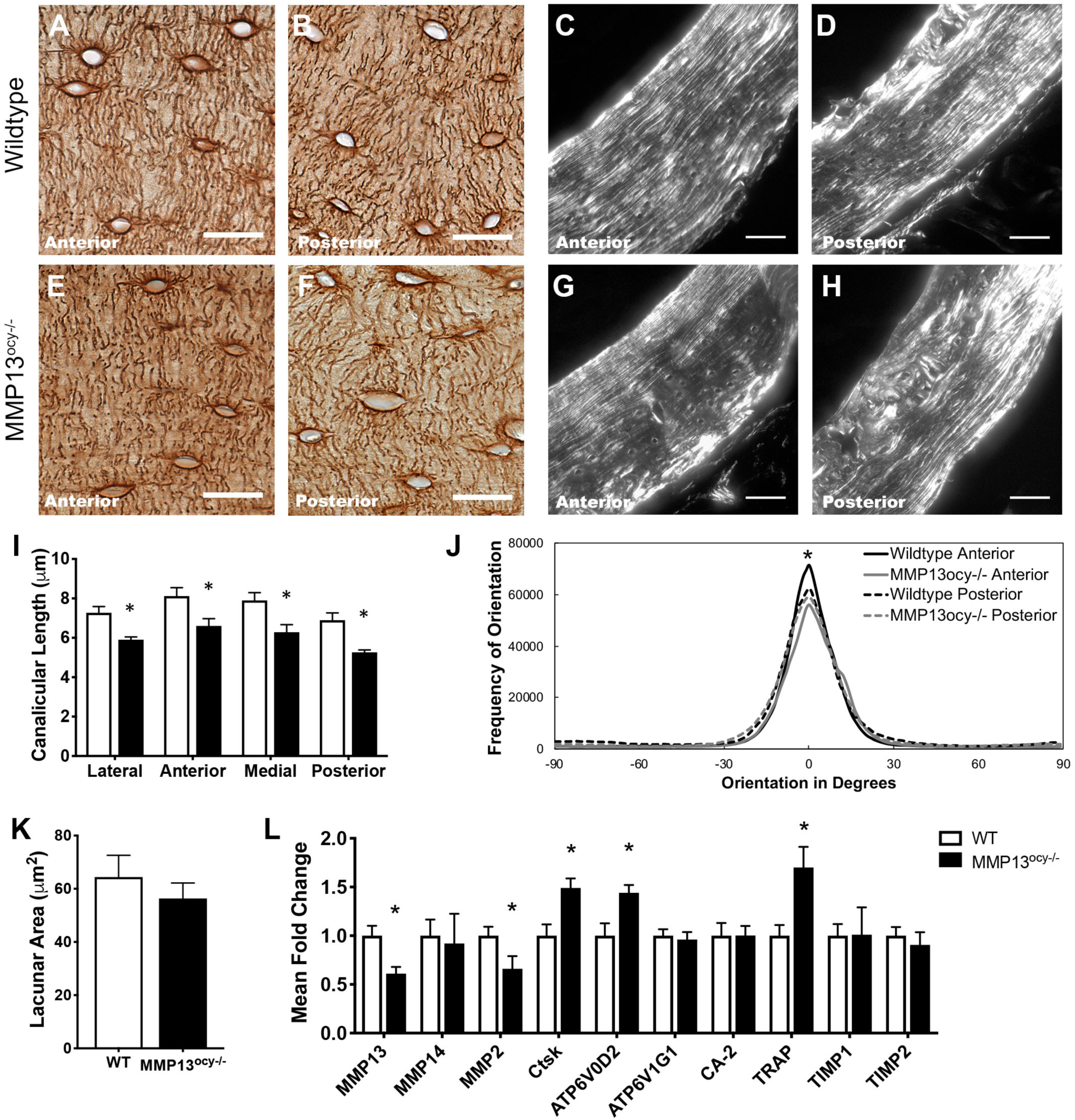
MMP13^ocy-/-^ cortical bone displays reductions in hallmarks of perilacunar/canalicular remodeling. Canalicular length in wildtype bone (A, B) was longer than that in MMP13^ocy-/-^ bone (E, F) in all regions sampled (I, n=7), while lacunar area was not statistically different (K, n=7). Collagen linearity was reduced in MMP13^ocy-/-^ bone (G, H, J, n=9) compared to wildtype bone (C, D, n=8) in the anterior quadrant (C, G), but not in other regions. qPCR of wildtype (n=8) and MMP13^ocy-/-^ (n=6) bone revealed both co-regulated and compensatory expression of PLR genes (L). Scale bars are 20 µm in A, B, E, F and 50 µm in C, D, G, H. *p<0.05 between genotypes by unpaired t-test.

The disruption of the LCN and ECM in MMP13^ocy-/-^ bone was accompanied by reduced expression of MMP2 and increased expression of Cathepsin K, ATP6V0D2, and TRAP (Fig. 5L). The increase in Cathepsin K is due to a change in osteocyte expression rather than osteoclast expression, which was verified by immunohistochemistry (Supp. Fig. 2C-D). These changes are consistent with prior observations of coordinated and compensatory changes in expression of genes required for matrix proteolysis and acid secretion in PLR (17,25).

Since changes to collagen, mineral, or LCN organization can affect bone biomechanical behavior (11,17,34), we tested mechanical properties of femurs from 2- and 4-month-old wildtype and MMP13^ocy-/-^ mice using 3-point bending. Small but significant decreases in whole-bone structural stiffness and ultimate load were detected in 4-month-old MMP13^ocy-/-^ bones (Table 1), consistent with minor deficiencies in both collagen and mineral (35,36). However, no significant changes were detected in yield properties, postyield displacement, or work-to-fracture, so cortical bone biomechanical outcomes were relatively insensitive to osteocyte-intrinsic MMP13 deficiency in this model. Overall, osteocyte-intrinsic MMP13 is required for PLR since its ablation disrupts the coordinated expression of PLR enzymes, the maintenance of canalicular networks and collagen organization, and bone mechanical properties.

**Table 1:** Flexural properties of wildtype and MMP13^ocy-/-^ femurs. Femurs from wildtype and MMP13^ocy-/-^ mice were broken with either anterior or posterior side in compression. In general, structural and tissue material properties were stronger in posterior (physiological) compression than anterior compression and get stronger with age. Ultimate load tended to be lower in MMP13^ocy-/-^ bones than wildtype bones, particularly in 4-month-old samples broken in anterior compression. MMP13^ocy-/-^ bones also had lower stiffness in this test configuration. *p<0.05 between genotypes, #p<0.065 between genotypes by unpaired t-test.

|  | Posterior compression, 2 months (n=9) |  | Posterior compression, 4 months (n=8-9) |  |
| --- | --- | --- | --- | --- |
|  | WT | MMP13<sup>ocy-/-</sup> | WT | MMP13<sup>ocy-/-</sup> |
| Bending Stiffness (N/mm) | 79.06 +/- 8.65 | 77.37 +/- 5.03 | 105.22 +/- 13.22 | 104.83 +/- 9.33 |
| Yield Load (N) | 11.57 +/- 1.55 | 10.94 +/- 1.63 | 13.05 +/- 2.43 | 13.86 +/- 1.55 |
| Ultimate Load (N) | 16.79 +/- 2.21 | 15.11 +/- 1.24 # | 19.02 +/- 2.08 | 18.24 +/- 1.63 |
| Postyield Displacement (mm) | 0.50 +/- 0.22 | 0.55 +/- 0.26 | 0.50 +/- 0.37 | 0.34 +/- 0.14 |
| Work to Fracture (N-mm) | 7.98 +/- 1.87 | 7.86 +/- 3.09 | 8.24 +/- 3.30 | 6.55 +/- 1.85 |
| Bending Modulus (GPa) | 10.37 +/- 1.52 | 11.75 +/- 2.70 | 14.35 +/- 1.08 | 14.72 +/- 3.07 |
| Yield Stress (MPa) | 158.04 +/- 11.51 | 167.55 +/- 37.78 | 186.41 +/- 35.28 | 200.00 +/- 38.15 |
|  | Anterior compression, 2 months (n=3-9) |  | Anterior compression, 4 months (n=8-9) |  |
|  | WT | MMP13<sup>ocy-/-</sup> | WT | MMP13<sup>ocy-/-</sup> |
| Bending Stiffness (N/mm) | 70.24 +/- 11.08 | 58.11 +/- 1.55 | 92.66 +/- 6.90 | 83.03 +/- 9.27 * |
| Yield Load (N) | 10.18 +/- 1.94 | 11.11 +/- 3.61 | 12.28 +/- 0.91 | 11.76 +/- 1.31 |
| Ultimate Load (N) | 13.25 +/- 1.65 | 13.61 +/- 1.67 | 17.12 +/- 1.34 | 15.64 +/- 1.35 * |
| Postyield Displacement (mm) | 0.91 +/- 0.53 | 0.95 +/- 0.40 | 0.28 +/- 0.10 | 0.34 +/- 0.19 |
| Work to Fracture (N-mm) | 8.91 +/- 2.32 | 9.16 +/- 2.40 | 5.09 +/- 1.50 | 5.27 +/- 2.01 |

### Increased cartilage degradation in MMP13^ocy-/-^ mice

Though subchondral bone clearly contributes to osteoarthritis, the responsible cellular mechanisms remain unclear (3,5,6). Given the strong association of PLR suppression with cartilage degeneration in human OA (Fig. 1-3), we tested the hypothesis that PLR suppression is sufficient to cause cartilage degeneration. Even with no differences in chondrocytic MMP13 expression (Supp. Fig. 2F-G, K-L), articular cartilage of 4-month-old wildtype (Fig. 6A-B) and MMP13^ocy-/-^ (Fig. 6D-E) mouse knees demonstrated clear histopathological differences. Relative to the smooth, proteoglycan-rich articular cartilage in wildtype knees, MMP13^ocy-/-^ cartilage had surface irregularities and depletion of proteoglycans. These characteristic features of degenerating articular cartilage were apparent in basal conditions on the medial and lateral tibial and femoral surfaces. Accordingly, using two established OA grading scales (37,38), MMP13^ocy-/-^ knees had statistically more cartilage degeneration than wildtype knees (Fig. 6C, F). Therefore, loss of MMP13 function in the subchondral bone osteocytes is sufficient to disrupt cartilage homeostasis, causing the appearance of osteoarthritic features in otherwise healthy cartilage. This result further suggests that the severe PLR suppression observed in OA human subchondral bone plays a causal role in the progression of joint disease.

**Figure 6:**
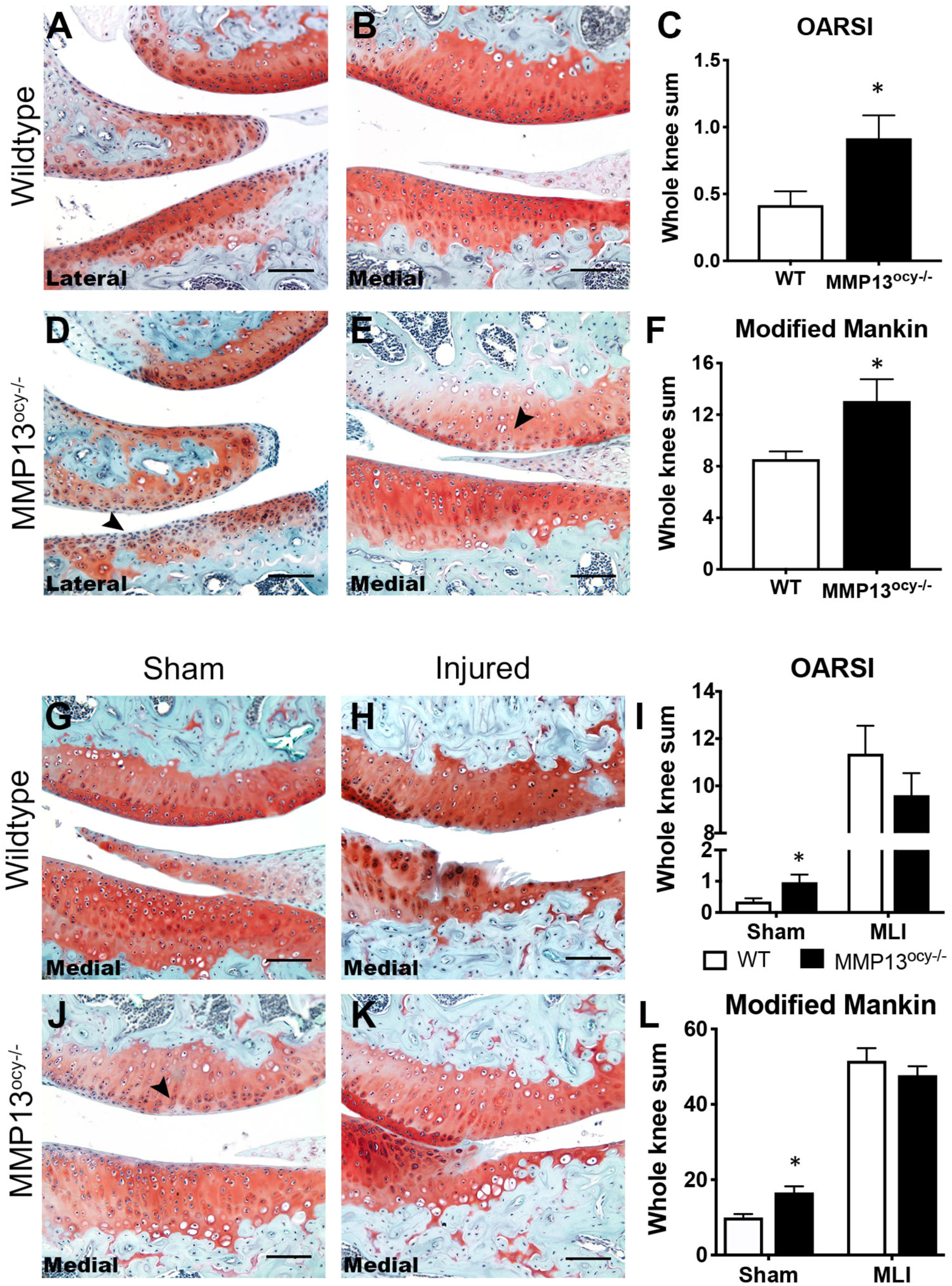
Knees of non-injured MMP13^ocy-/-^ mice show more cartilage damage than wildtype mice. Safranin-O stained joints of wildtype mice (A, B) show intact cartilage, while MMP13^ocy-/-^ mice (D, E) have areas of cartilage surface irregularity and proteoglycan depletion, leading to significantly higher OARSI (C) and modified Mankin (F) scores. In sham-injured joints (G, J), MMP13-dependent proteoglycan loss is still apparent, and meniscal-ligamentous injury (H, K) induces severe cartilage damage in both groups. On OARSI (I) and modified Mankin (L) grading scales, sham-injured MMP13^ocy-/-^ mice score significantly higher than wildtype mice (n=10), but scores are equivalent after injury (n=10 for wildtype, n=11 for MMP13^ocy-/-^). Arrows denote cartilage damage in non-injured joints. Scale bars are 100 µm. *p<0.05 between genotypes by unpaired t-test.

To assess the role of PLR suppression in post-traumatic cartilage degeneration, we used an established medial ligamentous injury (MLI) model to induce OA in wildtype and MMP13^ocy-/-^ knees (39). As in non-injured controls, sham-injured MMP13^ocy-/-^ joints showed more cartilage degeneration than their wildtype counterparts (Fig. 6G, J). Increases in chondrocyte hypertrophy and articular cartilage lesions, as well as proteoglycan loss, contributed to the more severe OA grade in sham-injured MMP13^ocy-/-^ knees (Fig. 6I, L). After injury, both genotypes experienced severe loss of articular cartilage and osteophyte formation (Fig. 6H, K), but no differences were found between genotypes. These data suggest that although MMP13-dependent osteocyte PLR is an important mechanism of healthy joint crosstalk, this role of PLR can be overshadowed by joint injury.

### Mechanisms of subchondral osteocyte influence on cartilage

To further understand why MMP13^ocy-/-^ mice are predisposed to cartilage degeneration, we evaluated biological and structural features of tibial subchondral bone in healthy and injured joints. Human osteoarthritic subchondral bone is characterized by sclerosis, collagen disorganization, and disrupted LCN (Fig. 1-3), so we hypothesized that these features would be increased in MMP13^ocy-/-^ subchondral bone. As in cortical bone (Fig. 5A, B, E, F), the canalicular length of osteocytes in MMP13^ocy-/-^ subchondral bone was visibly shorter than in wildtype bone (Fig. 7A, B, E, F). Canaliculi were 17-20% shorter in MMP13^ocy-/-^ subchondral bone compared to wildtype bone, and injured samples had 17-20% shorter canaliculi than in the corresponding sham-injured groups for each genotype (Fig. 7I). These findings highlight the bidirectional crosstalk in the joint, such that LCN disruption can cause cartilage degradation and that the LCN is sensitive to joint injury. Collagen organization in tibial subchondral bone was also disrupted by osteocytic MMP13 deficiency (Fig. 7C, G, J). Joint injury caused qualitatively less organized collagen (Fig. 7D, H), but no differences between genotypes were detected, Similarly, µCT of subchondral bone revealed a MMP13-dependent difference in bone volume fraction between the sham groups, but not between the injured groups (Fig. 7K). Thus, PLR suppression causes structural changes to subchondral bone that mimic joint injury, either of which can exacerbate cartilage degradation.

**Figure 7:**
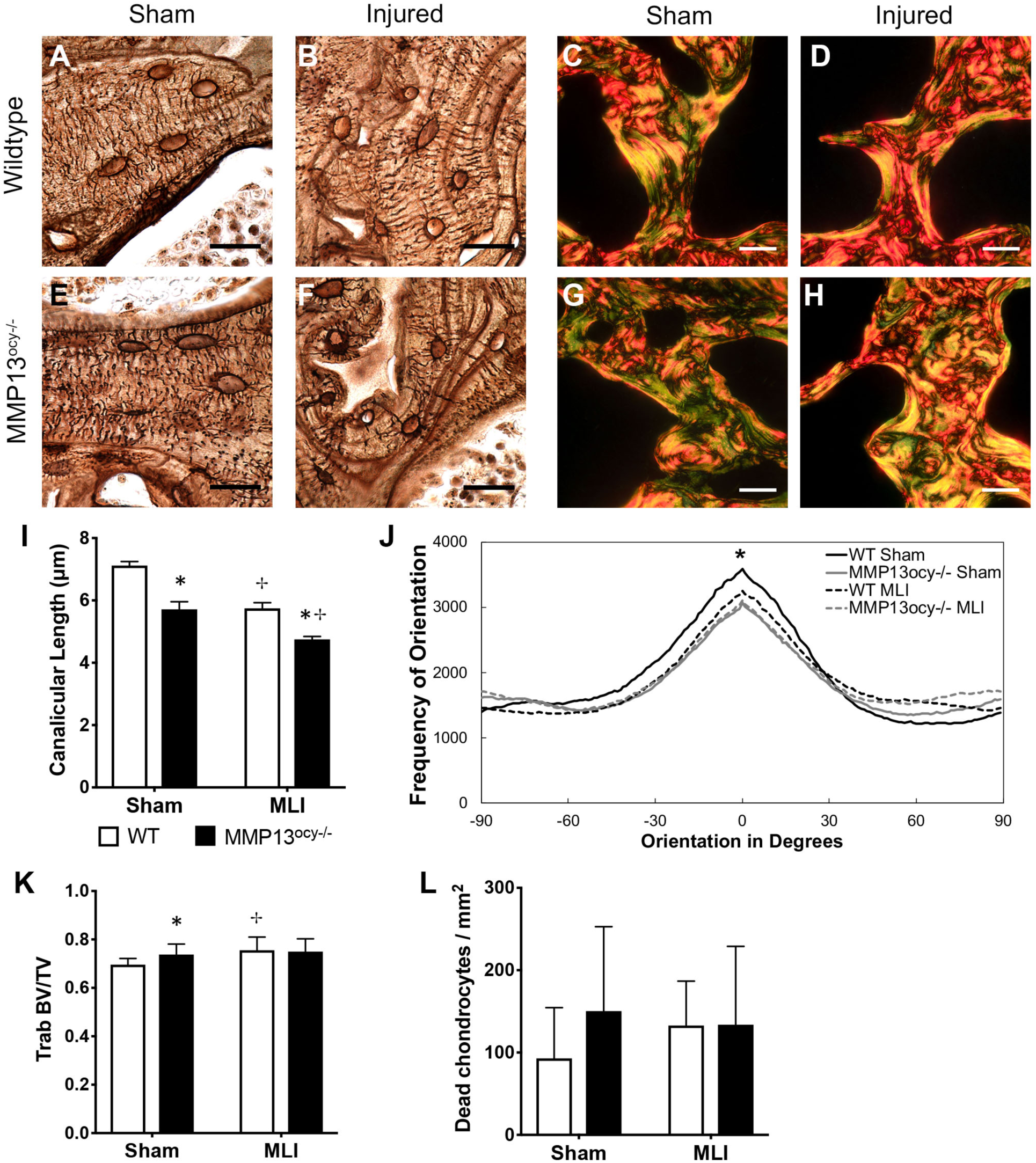
Subchondral bone shows MMP13-dependent changes before injury. Like in cortical bone, canalicular length in MMP13^ocy-/-^ subchondral tibias (E, F) is shorter than in wildtype subchondral tibias (A, B). The difference is statistically significant in injured and non-injured joints (I, n=6). Collagen organization is significantly affected by MMP13 in sham-injured subchondral tibia (C, G), but no difference is apparent after injury (D, H, J, n=6). Similarly, subchondral bone volume fraction is increased in sham-injured MMP13^ocy-/-^ tibias (n=10) compared to sham-injured wildtype tibias (n=10), but after injury wildtype (n=10) and MMP13^ocy-/-^ (n=11) joints are indistinguishable (K). No difference was detected in TUNEL-positive chondrocytes due to genotype or treatment (L, n=6). Scale bars are 20 µm in A, B, E, F and 50 µm in C, D, G, H. *p<0.05 between genotypes, +<0.05 between treatments by Holm-Sidak post-hoc tests.

Although a causal role for osteocytes in cartilage degeneration, to our knowledge, has not previously been demonstrated, there are multiple hypothetical mechanisms by which subchondral bone deterioration could exacerbate cartilage degeneration and drive OA (3,5,6). These include biological hypotheses such as cell death or vascular changes and mechanical hypotheses such as rod and plate distribution in subchondral bone (40–42). We tested the hypothesis that MMP13^ocy-/-^ bone causes increased chondrocyte catabolism and apoptosis. Using TUNEL staining, no significant difference in osteocyte or chondrocyte viability was detected due to osteocytic MMP13 deficiency or injury (Fig. 7L). While the lack of cell death after injury was surprising, cartilage area was significantly decreased by injury such that fewer chondrocytes could be counted. Likewise, differences in chondrocyte MMP13 expression were insufficient to explain the increased cartilage degeneration in MMP13^ocy-/-^ joints (Supp. Fig. 2O), though earlier timepoints may be required to observe these cellular responses (43,44). Additional studies of joint crosstalk will be needed to understand how biological mediators augment the structural changes in subchondral bone following PLR suppression to drive cartilage degeneration.

## DISCUSSION

This study advances our understanding of crosstalk between cartilage and subchondral bone by implicating osteocytes as causal drivers of joint disease in osteoarthritis. In both human and mouse joints, OA coincides with several outcomes of PLR suppression, including canalicular and collagen disorganization, hypermineralization, and repression of PLR enzyme expression. Furthermore, osteocyte-intrinsic deficiency in the key PLR enzyme MMP13 is sufficient to suppress PLR and induce premature osteoarthritis, with subchondral sclerosis and cartilage damage in otherwise healthy young mice. Using a surgical OA model, we find that cartilage crosstalk with osteocytes is bidirectional, since PLR is susceptible to injury even in wild-type mice. This study highlights a new and important role for osteocytes in the regulation of cartilage homeostasis and implicates PLR suppression as a causal mechanism in OA. Therefore, osteocytes emerge as a potential cellular target of new therapeutics to block or reverse OA progression.

A major gap in understanding OA is the identity of mechanisms responsible for the coupled degeneration of cartilage and subchondral bone. For example, human subchondral bone is sclerotic with altered osteocyte morphology beneath sites of cartilage deterioration (45,46). The current study advances the field by implicating osteocyte PLR as a cellular mechanism that couples bone and cartilage degeneration. These findings complement and extend beyond prior reports on altered osteocyte morphology, viability, and gene expression in human OA subchondral bone (45–49) by describing the profound effect of OA on the organizational pattern of the bone extracellular matrix, canalicular length, and lacunocanalicular network area. The OA-dependent change in LCN features is tightly correlated with a decrease in PLR enzyme expression (Fig. 3S, T). PLR is also suppressed in subchondral bone from patients with glucocorticoid-induced osteonecrosis and in a mouse model of glucocorticoid excess (25). Glucocorticoid-dependent repression of osteocyte enzyme expression in mice occurs within a week of treatment and precedes canalicular degeneration, collagen disorganization, and hypermineralization of subchondral bone (25), all of which mimic human osteonecrosis (23,24,50). While this study suggested a causal role for PLR suppression in joint disease, the well-known effects of glucocorticoids on many other cell types could not be excluded. The current findings, in which cartilage degeneration results from an osteocyte-intrinsic PLR defect, strengthen the idea that osteocytes play a causal role in joint disease.

To identify a causal role for PLR in cartilage degradation, we generated a novel mouse model of PLR repression using targeted deletion of MMP13 from osteocytes. We could not use the existing model of systemic MMP13 ablation because chondrocyte-derived MMP13 is one of the critical enzymes involved in cleavage of collagen II in OA (29), and several studies show the chondroprotective effect of MMP13-inhibition in chondrocytes (30,31). By allowing normal chondrocyte expression of MMP13 while reducing its expression in osteocytes, we observed evidence of osteocyte-dependent joint crosstalk, illustrating a novel cellular mechanism by which bone affects the overlying cartilage.

Several features of the bone phenotype in MMP13^ocy-/-^ mice resemble those in other models of MMP13 ablation and PLR suppression. Col1-Cre^+/-^;MMP13^fl/fl^ and systemic MMP13^-/-^ mouse models together implicate bone-derived MMP13 expression in trabecular bone remodeling, PLR, and bone quality (11,28,32). As in systemic MMP13^-/-^ bone, osteocyte-specific MMP13 ablation increased trabecular bone mass and disrupted canalicular and collagen organization. However, whereas systemic ablation caused decreased postyield deflection and decreased fracture toughness, we observed changes to cortical bone stiffness and ultimate strength only. This was surprising because we previously observed severe defects in bending modulus, yield stress, and toughness upon repression of PLR via osteocyte-specific inhibition of TGF-β signaling. Even with no change in cortical bone mass, work of fracture was reduced by 65% in TβRII^ocy-/-^ bone relative to wildtype controls (17). Possible explanations for these differences include the incomplete ablation of MMP13 in MMP13^ocy-/-^ bone, or the coordinated repression of multiple PLR enzymes in TβRII^ocy-/-^, but not in MMP13^ocy-/-^ bone. Whereas MMP2, MMP13, MMP14, Cathepsin K, and TRAP were repressed in TβRII^ocy-/-^ bone, MMP13^ocy-/-^ bone showed both co-repression (MMP2) and compensatory upregulation (Cathepsin K, TRAP) of PLR enzymes. Therefore, the partial reduction of PLR enzyme expression in the MMP13^ocy-/-^ model may be insufficient to substantially impact material properties and postyield behavior measured at the whole-bone scale. Overall this model was sufficient to suppress PLR in cortical and subchondral bone, consistent with the subchondral bone changes seen in end-stage human OA.

Much remains to be elucidated about the relative role of PLR in age-related joint degeneration and in post-traumatic OA (PTOA). Most animal models of OA involve a joint injury in which both bone and cartilage are affected, and which may additionally initiate inflammatory cascades (37,39,51–55), making it difficult to isolate the cell type responsible for cartilage degeneration. In our hands, MLI joint injury reduced canalicular length in both genotypes but did not affect MMP13^ocy-/-^ cartilage more severely. This suggests that joint injury can override preexisting defects in subchondral bone PLR (56). Review of additional timepoints post-injury, or use of a milder injury model, may provide a clearer illustration of how PLR contributes to PTOA pathogenesis. Basal phenotypes may be more representative of age-related joint degeneration, in which cartilage degeneration occurs without a known traumatic injury. The predisposition of mice with repressed PLR to develop cartilage wear is consistent with the idea that subchondral bone sclerosis leads to cartilage breakdown. For example, in a guinea pig model of spontaneous OA, trabecular rod loss and plate thickening precede significant cartilage degradation (40). Likewise, in non-human primates that spontaneously develop OA, subchondral bone thickening appears to precede cartilage fibrillation (57). In humans, longitudinal tracking of bone marrow lesions by MRI reveals the clinical relationship of subchondral changes to OA progression and knee pain (42). Although similar end-stage OA phenotypes can arise from age, injury, or other causes (58), PLR may play a causal role in some of these pathological mechanisms, but not others.

Unraveling the circumstances and mechanisms through which a bone-intrinsic defect in PLR exacerbates OA will require further studies. PLR has the potential to affect subchondral bone vascularity, microarchitecture, and mechanics, cartilage strain distribution, and cell-cell signaling, any of which could impact bone-cartilage crosstalk. The LCN, maintained by PLR, is important for solute transport, and transport between subchondral bone and cartilage is altered in mouse and human OA (19,59,60). Changes in subchondral bone volume fraction (41,61) and geometry (40), similar to those seen in MMP13^ocy-/-^ bone, can precede cartilage pathology, possibly by changing strain distribution in the overlying cartilage. Biological mechanisms might include the action on chondrocytes of factors secreted by osteocytes or released from the bone ECM during PLR (62,63). Furthermore, this is likely not a unidirectional process. In fact, LCN measurements between medial and lateral compartments in humans and between injured and non-injured mice suggest that cartilage injury can cause further repression of PLR in subchondral bone, possibly forming a deleterious feedback loop. If the critical points at which PLR causes and responds to cartilage degradation can be identified, osteocytes may emerge as a potential cellular target of new therapeutics to prevent or treat OA. Although many questions remain about the role and regulation of PLR in OA, our data collectively suggests that osteocyte PLR dynamically maintains subchondral bone homeostasis and joint crosstalk, and that its disruption can exacerbate joint disease.

## MATERIALS & METHODS

### Human donor population and specimen preparation

Five male subjects with clinically diagnosed stage IV osteoarthritis of the tibial plateau, who were scheduled for total knee arthroplasty, were recruited for this study (Supp. Fig. 1A). Recruitment occurred through referral from orthopedic surgeons at a Department of Veterans Affairs Medical Center. All samples were collected from patients with OA of the femorotibial joint as described in protocols that were reviewed and approved by our Human Subjects Protection Program Institutional Review Board. Informed consent was obtained from each study participant prior to enrollment. Five freshly harvested cadaveric human tibial plateaus from age- and gender-matched donors without history of OA, osteonecrosis, osteoporosis, or fractures were collected through the Willed Body Program at University of California, San Francisco for use as controls. Patients with OA and healthy cadaveric controls had similar BMI.

Each tibial plateau was removed *en bloc* (Supp. Fig. 1C-D), and X-rays were collected to evaluate the severity of subchondral bone deterioration (Supp. Fig. 1G-I). To facilitate comparison of the subchondral bone between the lateral and medial side of the joint, which was more severely affected by OA in these samples, each specimen was cut into 8-10 mm thick coronal slabs with a band saw (Supp. Fig. 1E, F). Subchondral bone was compared between the medial and lateral regions of interest on the same osteoarthritic tibial plateau, as well as with comparable regions of cadaveric tibial plateaus. Data was collected from five samples per group for all outcomes unless otherwise specified in figure legends.

### Mice

To test the role of osteocytic MMP13 in bone and joint health, we generated mice with osteocyte-specific ablation of MMP13. Homozygous MMP13^fl/fl^ mice have loxP sites flanking exons 3, 4, and 5, which encode the enzyme’s active site (32). Hemizygous DMP1-Cre^+\-^ mice (9.6-kb promoter) express Cre in osteocytes as well as in odontoblasts, with little to no expression in osteoblasts, muscle, kidney, liver, or brain (33). Mice were bred at UCSF to generate wildtype (DMP1-Cre^-/-^; MMP13^fl/fl^) and MMP13^ocy-/-^ (DMP1-Cre^+/-^; MMP13^fl/fl^) mice. For this study, only male mice were utilized. The procedures for animal experiments were approved by the Institutional Animal Care and Use Committee at the University of California, San Francisco. Six to eleven biological replicates were used for each outcome, with exact n given in figure legends.

### Histology

Human tibial plateaus were fixed in 10% neutral buffered formalin (NBF) and incubated in 10% di- and tetra-sodium EDTA for 56-60 days until fully decalcified, or in an Ion Exchange Decalcification Unit (American Master Technologies) for 5-6 days, followed by serial ethanol dehydrations and paraffin embedding. Paraffin sections (7 µm thick) in the coronal plane were generated for polarized light microscopy, Safranin-O with Fast Green stain, Ploton silver stain, and immunohistochemistry. To standardize evaluation, a consistent region of subchondral bone was selected for evaluation in the medial and lateral areas of each specimen (Fig. 1A-B). For each specimen, values were collected from 5 high-powered field images per region of interest. Within each region, these values were averaged to obtain a mean value for each specimen. Each quantitative average represents an average across all specimens.

Intact mouse knee joints and proximal femurs were fixed in 10% NBF and decalcified for two weeks in EDTA, followed by serial ethanol dehydration and paraffin embedding. Knees were embedded at 90 degrees of flexion and sectioned in the frontal plane. Femora were embedded and sectioned axially to generate 6 µm sections All brightfield imaging was conducted on a Nikon Eclipse E800 microscope.

#### Safranin-O/ Fast Green stain and OA scoring

Safranin-O with Fast Green was used to visualize the cartilage quality of the tibial plateaus using a protocol adapted from University of Rochester (64). Briefly, sections were deparaffinized, rehydrated, and incubated in Weigert’s Iron Hematoxylin for 3 minutes. Stained slides were then washed in water and differentiated in 1% Acid-Alcohol for 15 seconds. Slides were then stained with a 0.02% aqueous Fast Green solution for 5 minutes and differentiated with 1% acetic acid for 30 seconds. Slides were then washed with water and stained in a 1% Safranin-O solution for 10 minutes and subsequently dehydrated, cleared, and mounted.

For human tibial plateaus, standardized OARSI grading (65,66) was used to assess OA in Safranin-O stained histological sections by two orthopedic surgeons. For murine samples, Safranin-O staining was conducted on sections of the knee in a plane where the ACL and PCL were visible to maintain constant region of interest. Each quadrant of the knee (medial and lateral tibia and femur) was graded by three blinded graders using OARSI (38) and modified Mankin (37) scales. For each sample, the numerical scores of all graders were averaged to obtain a mean score. Mean scores were then averaged within each group.

#### Analysis of collagen orientation by picrosirius red stain

Polarized light microscopy was performed on deparaffinized sections stained in a saturated aqueous solution of picric acid with 0.1% Direct Red-80, as described (28,67) to visualize collagen fibril orientation. During microscopy, polarized filters were rotated to achieve the maximum birefringence before capturing each image. Red channel images were processed using the OrientationJ plugin for ImageJ as described (68). Statistical analysis was performed as described below.

#### Analysis of the lacunocanalicular network by Ploton silver stain

To visualize the osteocyte lacunocanalicular network, sections were deparaffinized and incubated in two-parts 50% silver nitrate and one-part 1% formic acid in 2% gelatin solution for 55 minutes, as described (28,69). Stained slides were then washed in 5% sodium thiosulfate for 10 minutes and subsequently dehydrated, cleared, and mounted. Images were acquired at 100x for analysis. For human tibial plateau subchondral bone, quantification of the lacunocanalicular area was performed with ImageJ by thresholding gray-scale images for dark, silver stained lacunae and canaliculi. The resulting area was normalized to total bone area in each image captured. Canalicular length was analyzed with ImageJ by individually measuring canaliculi surrounding osteocytes (average 10 canaliculi per osteocyte). For murine cortical bone, two images were acquired in each quadrant of an axial section (medial, lateral, anterior and posterior). In murine tibial plateau subchondral bone, four images were captured on the medial and lateral side of the joint. In each image, 10 canaliculi on 3 osteocytes were traced using ImageJ to calculate mean canalicular length. Lacunar area was calculated with a custom, commercially available StrataQuest application (TissueGnostics).

#### Analysis of PLR enzyme expression by immunohistochemistry (IHC)

IHC was used to examine protein localization qualitatively and semi-quantitatively (i.e. % positively stained cells). For IHC, slides were deparaffinized and hydrated prior to incubation in Innovex Uni-Trieve low temperature retrieval solution (NB325) in a 40°C water bath for 24 hours (human) or in a 65 °C water bath for 30 minutes (mouse). Endogenous peroxidase activity was quenched using 3% H_2_O_2_ for 10 minutes at room temperature. For the subsequent steps, Innovex Universal Animal Immunohistochemistry Kit (329ANK) was utilized. Samples were blocked with Fc-Block and Background Buster for 45 minutes each at room temperature. Primary antibodies were diluted in PBS (anti-MMP13, 1:100, Abcam 39012; anti-CatK, 1:50, Abcam 19027) and incubated in a humid chamber at 4°C for 24 hours. Secondary linking antibody and HRP-enzyme were both used at room temperature for 10 minutes each. Fresh DAB solution was applied and incubated at room temperature for 5 minutes prior to washing with tap water and mounting with Innovex Advantage aqueous mounting medium (NB300). Negative controls were performed by substituting Innovex rabbit negative control sera in place of primary antibody. Quantification was performed with the help of ImageJ cell counter plug-in to determine the average percentage of MMP13-positive and CatK-positive osteocytes, relative to the total number of osteocytes in each 40x magnified visual field.

In murine cortical bone, MMP13 expression was additionally visualized and quantified using immunofluorescence. Sections were deparaffinized and antigens were retrieved with Uni-Trieve solution as above. Sections were blocked with Background Buster (Innovex) for 10 minutes, incubated with PBS/0.1% Tween for 5 minutes, and then incubated overnight with rabbit polyclonal anti-MMP13 antibody (1:50, ab39012). After washes in PBS, secondary goat-anti-rabbit antibody (1:1000, ab150080) was applied for 60 minutes, background was reduced with copper sulfate for 10 minutes, and slides were mounted with Prolong Gold antifade reagent with DAPI. Images were acquired on a Leica DMi8 confocal microscope. The percentage of MMP13-expressing osteocytes was calculated relative to the total number of osteocytes in at least two 40x fields per sample for 8-9 mice per genotype.

#### Analysis of cell death by TUNEL assay

To detect osteocyte and chondrocyte death, mouse knee sections were deparaffinized and permeabilized in 0.1% sodium citrate with 0.1% Triton X-100 for 8 minutes. For a positive control, two sections were treated with DNase for 10 minutes at room temperature to induce DNA strand breaks. Then all samples were incubated with TUNEL reaction mix for 60 minutes at 37 °C (Roche). After washing, slides were mounted with Prolong Gold antifade reagent with DAPI and imaged on a Leica DMi8 confocal microscope. Total number of labeled osteocytes and chondrocytes per bone or cartilage area was calculated in the medial and lateral compartments of the tibia for six knees per group.

### Synchrotron Radiation X-ray Computed Micro-Tomography (SRµT)

To visualize and quantify bone mineralization, 4-mm-wide specimens of cartilage and subchondral bone were imaged by SRµT at beamline 8.3.2 of the Advanced Light Source (ALS) (Lawrence Berkeley National Laboratory, Berkeley) as described (25). Briefly, transmission radiographs were taken over a 180° rotation with a monochromatic energy of 20 keV and an exposure time of 800 ms. Computational reconstruction of 3D images reveals bone microstructure at 1.3 µm/pixel (5X lens, LuAG scintillator). Images were segmented using ImageJ by binarization of the bone volume morphology. 3D visualization and quantification of bone mineral density was performed using Avizo (Visualization Sciences Group). Data was collected from n=2 for control medial, n=2 for control lateral, n=4 for OA medial, and n=5 for OA lateral human tibial plateaus. Statistical analysis was performed as described below.

### Micro-computed tomography (μCT)

For skeletal phenotyping, femurs were harvested from male mice at 13 weeks old and stored in 70% ethanol. Cortical analysis was conducted in a 1-mm region equidistant from the proximal and distal ends of the bone. Trabecular analysis was conducted in a 2-mm region immediately proximal to the distal growth plate. For subchondral bone analyses, knee joints were harvested from 16-week-old males and stored in saline solution at −20° C. A 4-mm region centered on the joint was scanned. Medial and lateral tibial subchondral bone were delineated 200 µm from the proximal surface of the tibia and extending for 250 µm distally. The medial and lateral femoral condyles were designated 200 µm from the distal end of the femur and extended proximally 200 µm. All samples were scanned using a Scanco µCT50 specimen scanner with an X-ray potential of 55 kVp, current of 109 µA, and voxel size of 10 µm. Thresholding and quantification was performed as previously described (17,25).

### Quantitative RT-PCR analysis

Humeri from wildtype and MMP13^ocy-/-^ mice were cleaned of muscle and periosteum, epiphyses were trimmed, and marrow was removed by centrifugation. Bones were snap-frozen in liquid nitrogen prior to homogenization in TRIzol (Invitrogen), as described (17,25). Briefly, mRNA was purified using the RNeasy Mini Kit following manufacturer’s instructions (Qiagen). RNA was reverse-transcribed using the iScript cDNA synthesis kit. qPCR was performed using Taqman probes for β-actin (assay #Mm02619580_g1) and MMP13 (assay #Mm00439491_m1 which targets exons 4-5) and using iQ SYBR Green Supermix (BioRad) with β-actin as the housekeeping gene (primer sequences given in Supp. Table 1). Expression was then quantified by the ΔΔCt method (70).

### Flexural strength tests

Whole-bone biomechanical properties were measured in femurs isolated from 2-month and 4-month-old wildtype and MMP13^ocy-/-^ mice. Whole hydrated femurs were loaded to failure in 3-point bending using a Bose Electroforce 3200 test frame. One femur per mouse was broken in the direction of primary physiological bending (posterior compression) and the other was broken against the direction of physiological bending (anterior compression). An 8-mm span was chosen because it was approximately 50% of the bone length. Tests were performed in air at a fixed displacement rate of 10 µm/s. Whole-bone stiffness was calculated from the linear portion of the load-displacement curve and yield was designated as the point where a line representing a 10% loss in stiffness intersected the load-displacement curve (71). Following fracture, bone cross sections were imaged by scanning electron microscopy on a Sigma 500 VP FE-SEM (Zeiss) at an excitation voltage of 15 kV and a partial pressure of 35 Pa. Measurements of cross-sectional diameter and thickness were acquired in ImageJ and used to calculate moment of inertia assuming an elliptical cross section. These geometric parameters were used to convert the load-displacement data to stress-strain data in order to measure tissue modulus, tissue stress and tissue strain with standard beam theory equations (72).

### MLI surgery

8-week-old male mice were separated into three groups: control, sham, and meniscal-ligamentous injury (MLI) (39). Under general isofluorane anesthesia, both hind limbs of MLI animals were shaved and sterilized. A bilateral approach was chosen in order to minimize effects of altered biomechanics arising from a single knee injury, as previously described (73,74). Briefly, medial incisions through the skin and joint capsule were made adjacent to the patella to expose the medial collateral ligament, which was transected. The medial meniscus was then removed. Sham-injured animals received bilateral incisions without MCL transection or meniscus dissection. Skin incisions were closed with sutures and animals received an injection of long-acting buprenorphine analgesic. Control animals did not receive anesthesia or analgesics. All animals were allowed unrestricted activity, food, and water. At 16 weeks of age, animals were euthanized and hind limbs collected for histological and radiographic analyses.

### Statistics

Values measured from histological analysis are expressed as mean +/- SEM, and other types of data are expressed as mean +/- SD. Comparisons between two groups were tested with unpaired two-tailed Student’s t-test. Comparisons between disease state and region in human specimens or between genotype and injury in mice were tested with two-way ANOVA followed by Holm-Sidak post-hoc tests. The Clinical and Translational Science Institute Statistical Consulting Service at UCSF evaluated significant differences in the distributions of human collagen organization and mineral density (SRµT) among each group using a mixed model with random intercepts. A linear model was used for the fixed effects, and the outcome was logarithmically transformed. In all figures, p-values less than 0.05 were considered statistically significant and are reported as such.

### Study Approval

The Human Subjects Protection Program Institutional Review Board at UCSF and the Research Committee at the San Francisco Veterans Affairs Medical Center reviewed and approved the protocols for human sample collection. Written informed consent was obtained from each study participant prior to enrollment.

## Author Contributions

Study design: all authors. Study conduct: CM, JW, CY, AF, CA, KB, AK and TF. Data collection: CM, JW, CA and KB. Data analysis: CM, JW, CA, KB, AK, TV and TA. Data interpretation: CM, JW, JL, AD, AK, TV and TA. Drafting manuscript: CM, JW and TA. Revising manuscript content: all authors. Reading and approving of final version of manuscript: all authors. TA takes responsibility for the integrity of the data analysis.

## Acknowledgements

This research was supported by the NIH-NIDCR grant R01 DE019284 (T.A.), the Department of Defense (DoD) grant PRORP OR130191 (T.A.), the Read Research Foundation (T.A.), NSF GRFP 1650113 (C.M.), NSF CDMI (T.A.), and the NIH-NIAMS grant P30 AR066262-01 (J.L., T.A.). The authors gratefully acknowledge Ellen Liebenberg for expert histological assistance, and Jennifer Salinas and Leah Lorget for assistance with histological analysis. The authors acknowledge the use of x-ray synchrotron beamlines 8.3.2 at the Advanced Light Source at the Lawrence Berkeley National Laboratory.

---

**Supplemental Figure 1:**
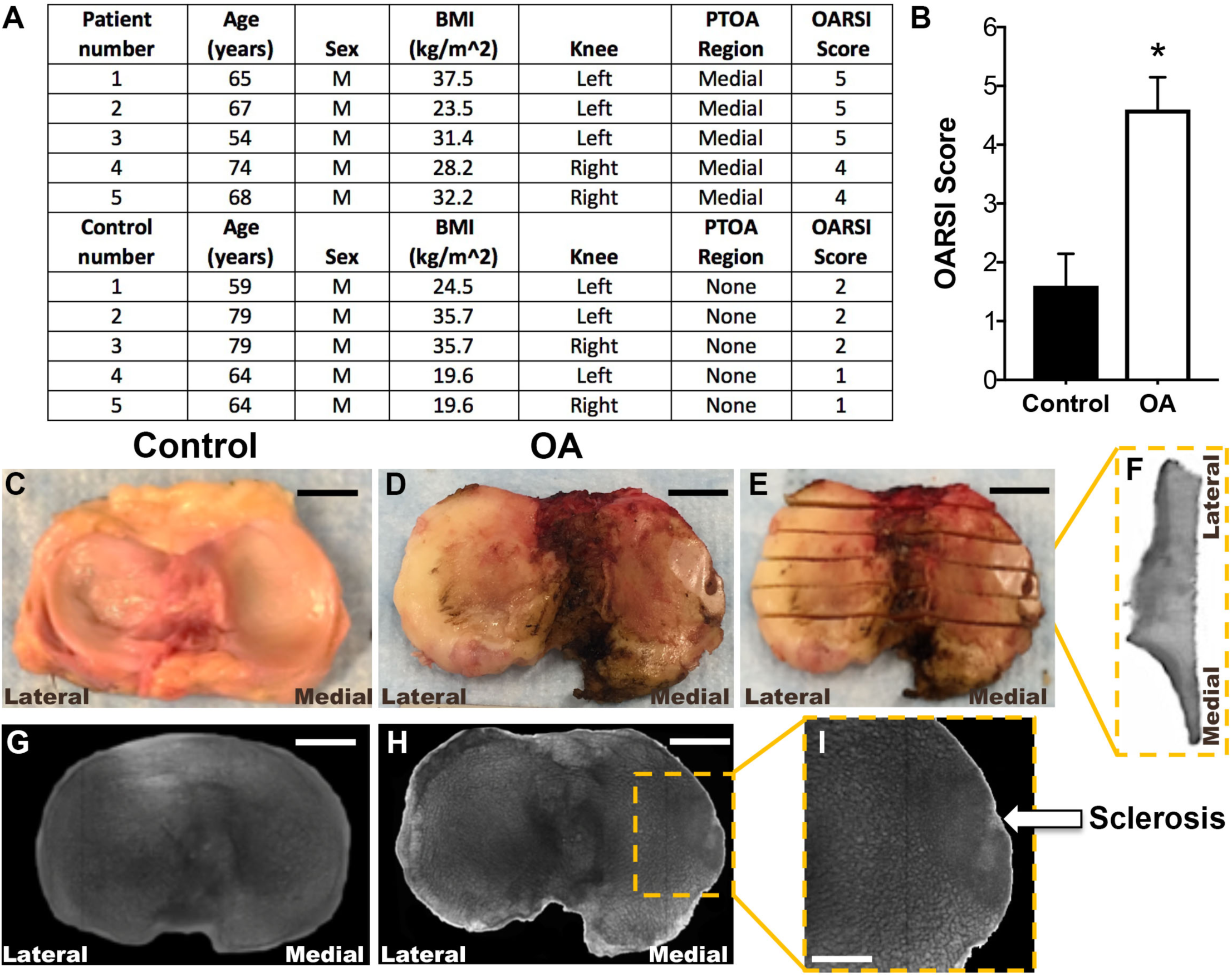
Characteristics of human tibial plateau specimens. Human tibial plateau specimens from patients with medial osteoarthritis undergoing total knee replacement surgery were compared to control cadaveric human tibial plateaus from donors with no history of osteoarthritis, osteonecrosis, osteoporosis, or fractures. Donations were controlled for age, sex, and body mass index (BMI) (A). OARSI scores (B) were consistent with gross (C, D) and radiographic (G, H) evidence of OA in surgical specimens. Gross cartilage degradation and sclerosis predominated in the medial compartment of OA specimens (I). For further histological analysis, intact specimens were cut into six 8-10 mm thick coronal slabs (E, F) Scale bars are 2 cm in C-H and 1 cm in I. *p<0.05 between conditions by unpaired t-test.

**Supplemental Figure 2:**
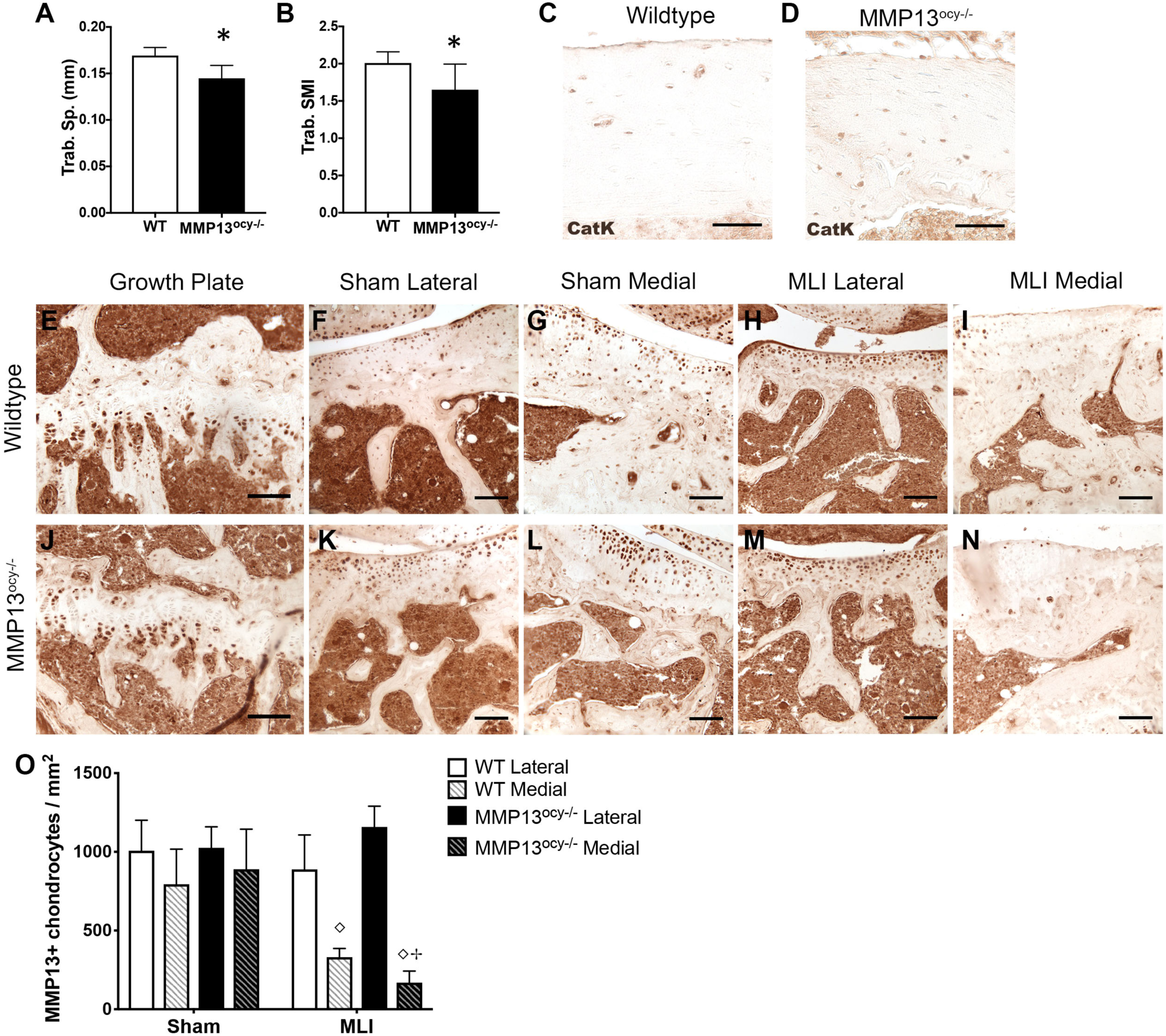
MMP13 expression in growth plate and articular cartilage chondrocytes is unaffected by deletion from osteocytes. MMP13^ocy-/-^ bones have increased bone volume fraction accompanied by a decrease in trabecular spacing (A) and in trabecular SMI (B). Immunohistochemistry for Cathepsin K reveals increased osteocyte expression in MMP13^ocy-/-^ cortical bones (C, D). As expected, the number of MMP13-expressing osteocytes in subchondral bone is lower in MMP13^ocy-/-^ mice than in wildtype mice, but no further differences were apparent after injury (F-I, K-N). MMP13 expression in articular cartilage and growth plate chondrocytes is not significantly different between genotypes (E-O). Consistent with prior reports of end-stage OA (44), we observed a stark loss of MMP13 expression in chondrocytes on the medial side of injured joints (I, N) compared to the lateral side and compared to medial chondrocytes in the corresponding sham groups. Scale bars are 50 µm in C, D and 100 µm in E-N. *p<0.05 between genotypes by unpaired t-test. ◊p<0.05 between regions, +<0.05 between treatments by Holm-Sidak post-hoc tests.

**Supplemental Table 1:**
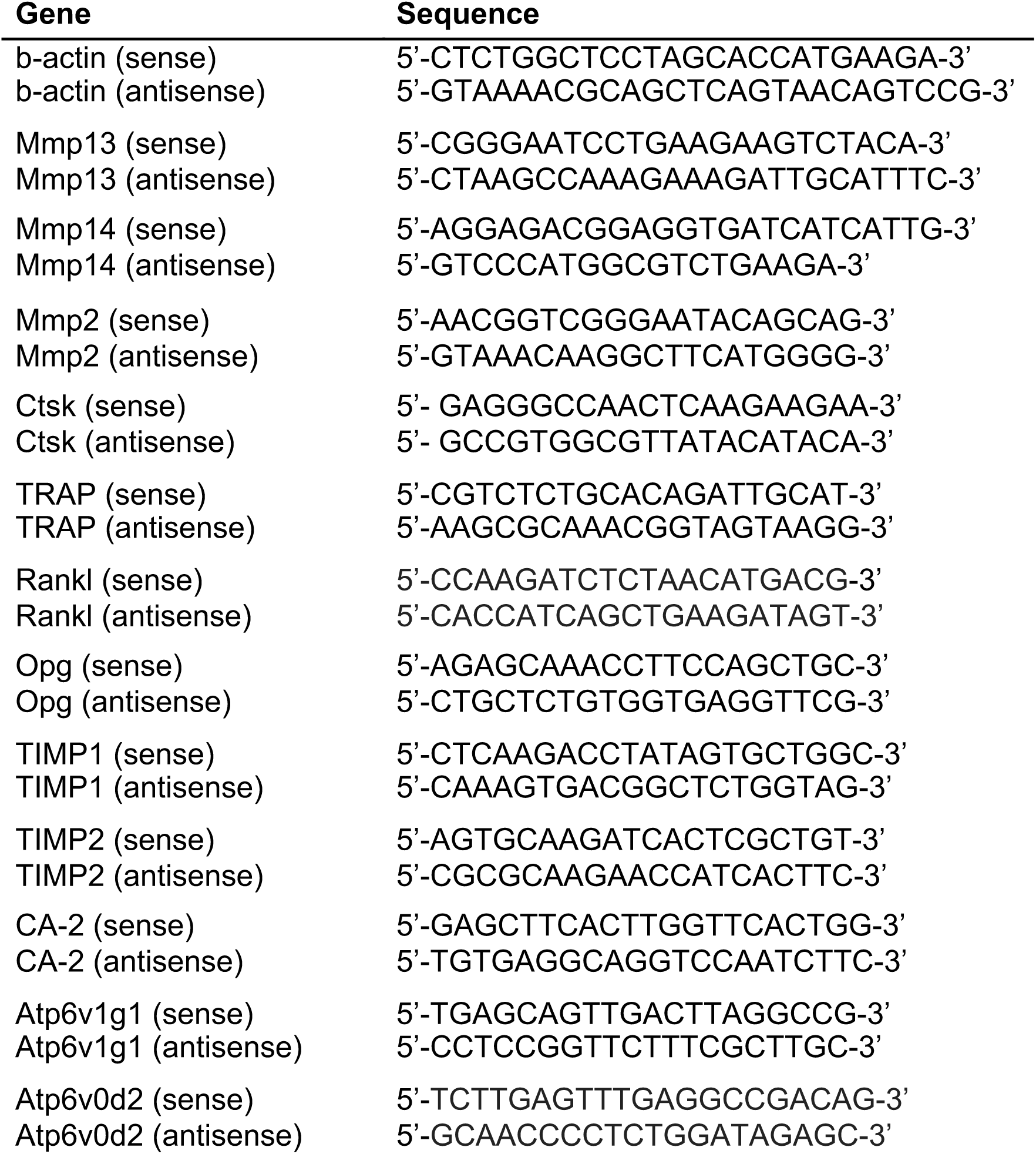
SYBR primer sequences used for gene expression analysis of murine mRNA.

## Notes

**Conflict of interest statement** The authors have declared that no conflict of interest exists.

